## Supplemental Figures for "Single Cell Analysis Reveals Immune Cell-Adipocyte Crosstalk Regulating the Transcription of Thermogenic Adipocytes"

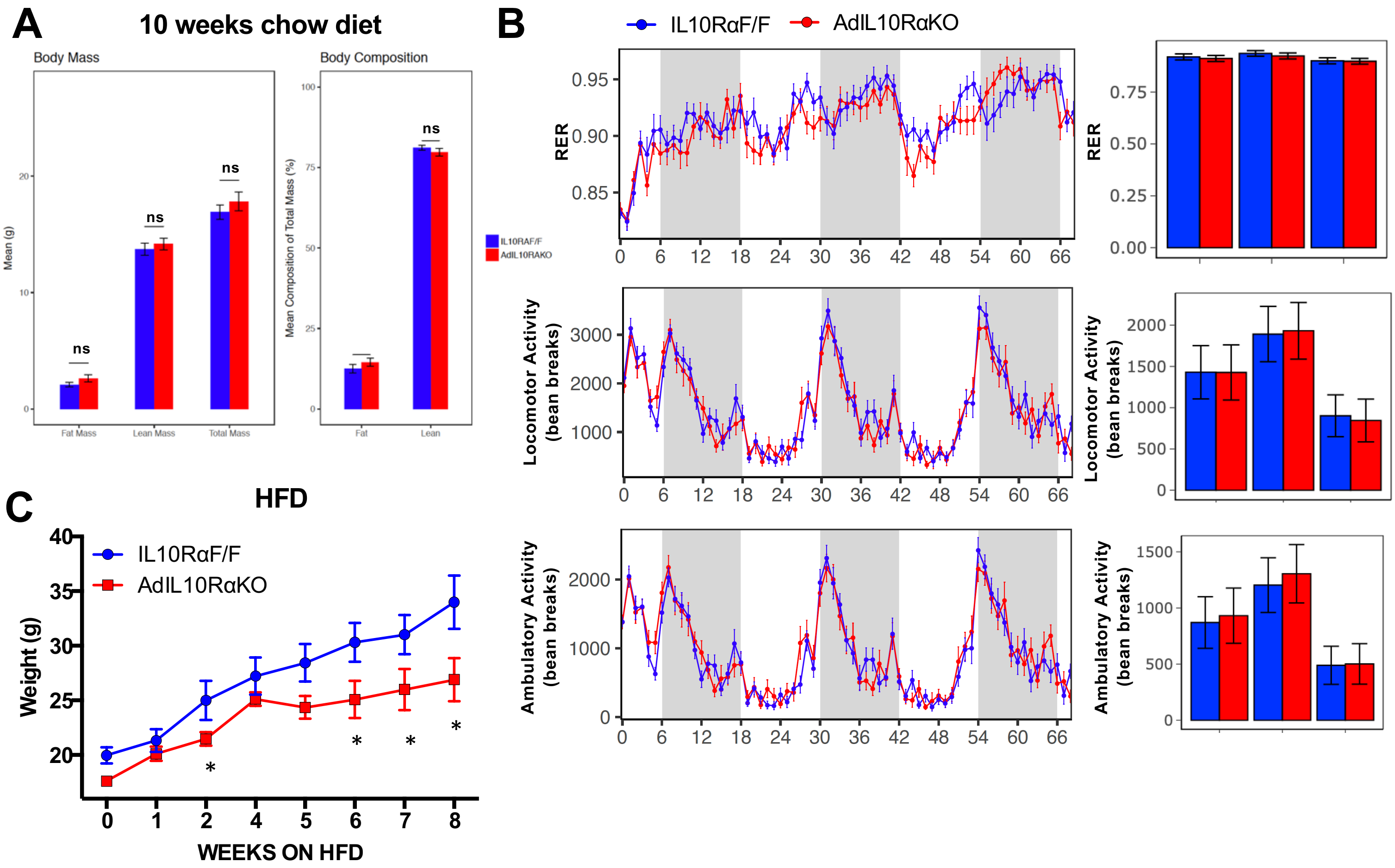


D

| GO biological process complete | # in Mouse Genome | # in Input Gene List | # Genes Expected | Fold Enrichment | P value |
|---|---|---|---|---|---|
| positive regulation of lipid biosynthetic process | 92 | 8 | 0.66 | 12.18 | 4.93E-03 |
| --> positive regulation of lipid metabolic process | 153 | 11 | 1.09 | 10.07 | 2.24E-04 |
| --> positive regulation of biological process | 5893 | 70 | 42.09 | 1.66 | 1.81E-02 |
| --> regulation of metabolic process | 5649 | 69 | 40.35 | 1.71 | 7.84E-03 |
| --> regulation of lipid metabolic process | 328 | 14 | 2.34 | 5.98 | 1.40E-03 |
| --> regulation of lipid biosynthetic process | 167 | 11 | 1.19 | 9.22 | 5.20E-04 |
| positive regulation of cold-induced thermogenesis | 95 | 8 | 0.68 | 11.79 | 6.19E-03 |
| --> regulation of cold-induced thermogenesis | 141 | 9 | 1.01 | 8.94 | 1.10E-02 |
| regulation of lipid transport | 112 | 8 | 0.8 | 10 | 1.98E-02 |
| --> regulation of lipid localization | 142 | 10 | 1.01 | 9.86 | 1.18E-03 |
| lipid localization | 278 | 12 | 1.99 | 6.04 | 9.87E-03 |
| lipid metabolic process | 1045 | 26 | 7.46 | 3.48 | 3.21E-04 |
| --> metabolic process | 7138 | 79 | 50.98 | 1.55 | 4.30E-02 |
| regulation of apoptotic process | 1468 | 31 | 10.48 | 2.96 | 5.20E-04 |
| --> regulation of programmed cell death | 1488 | 31 | 10.63 | 2.92 | 7.01E-04 |
| --> regulation of cell death | 1636 | 35 | 11.68 | 3 | 4.33E-05 |
| regulation of catalytic activity | 1793 | 32 | 12.81 | 2.5 | 1.24E-02 |
| regulation of signal transduction | 2704 | 45 | 19.31 | 2.33 | 5.28E-04 |
| --> regulation of cell communication | 3110 | 51 | 22.21 | 2.3 | 5.51E-05 |
| --> regulation of signaling | 3125 | 51 | 22.32 | 2.29 | 6.28E-05 |
| --> regulation of response to stimulus | 3795 | 54 | 27.1 | 1.99 | 2.09E-03 |
| negative regulation of cellular process | 4443 | 57 | 31.73 | 1.8 | 2.98E-02 |
| --> negative regulation of biological process | 4959 | 61 | 35.42 | 1.72 | 4.62E-02 |
| Unclassified | 1822 | 3 | 13.01 | 0.23 | 0.00E+00 |

| | # in Mouse Genome | # in Input Gene List | # Genes Expected | Fold Enrichment | P value |
|---|---|---|---|---|---|
| adaptive immune response | 468 | 9 | 0.97 | 9.31 | 4.07E-03 |
| Unclassified | 1822 | 3 | 3.76 | 0.8 | 0.00E+00 |

**Rajbhandari Fig. S1**

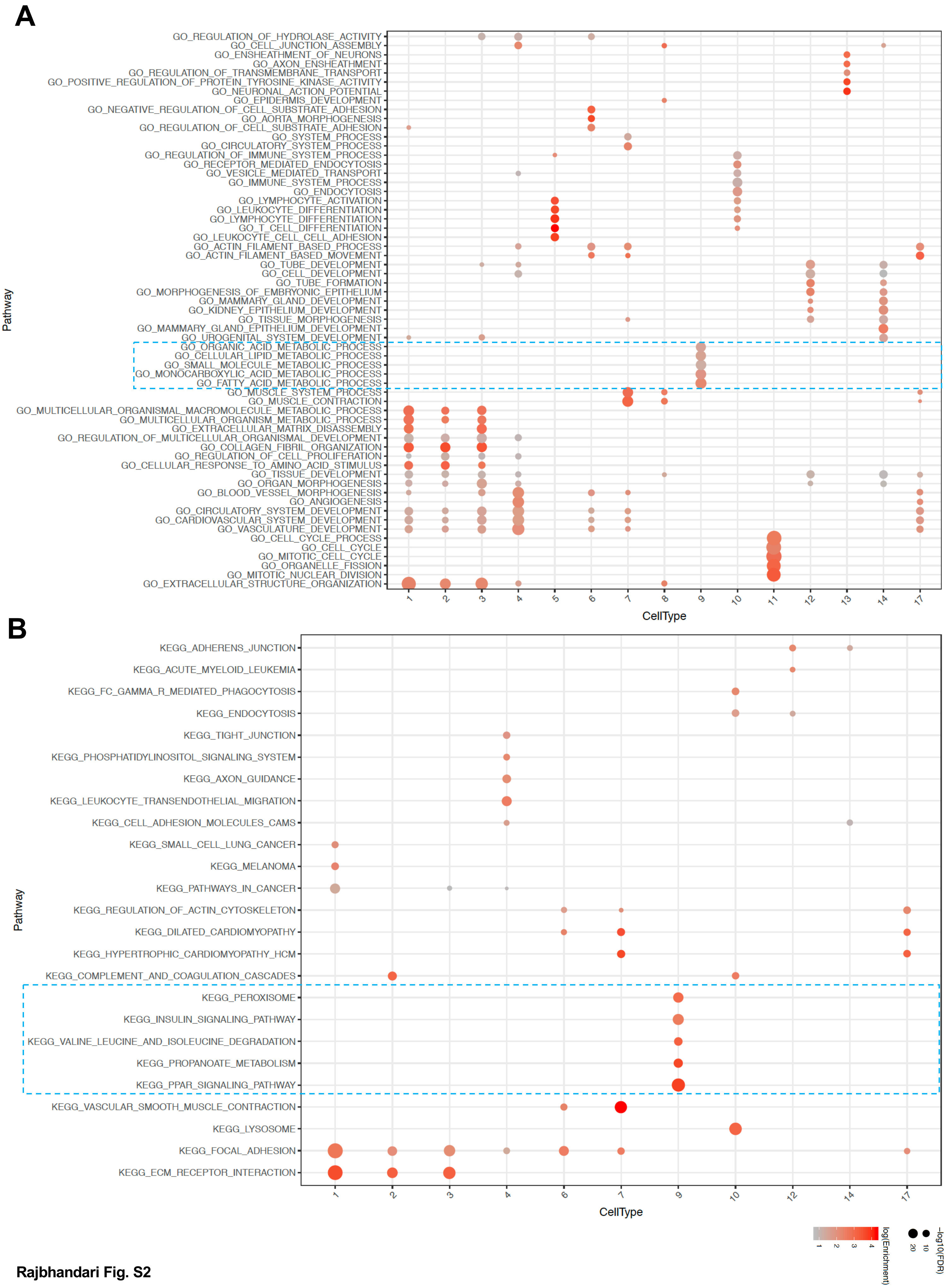


Rajbhandari Fig. S2

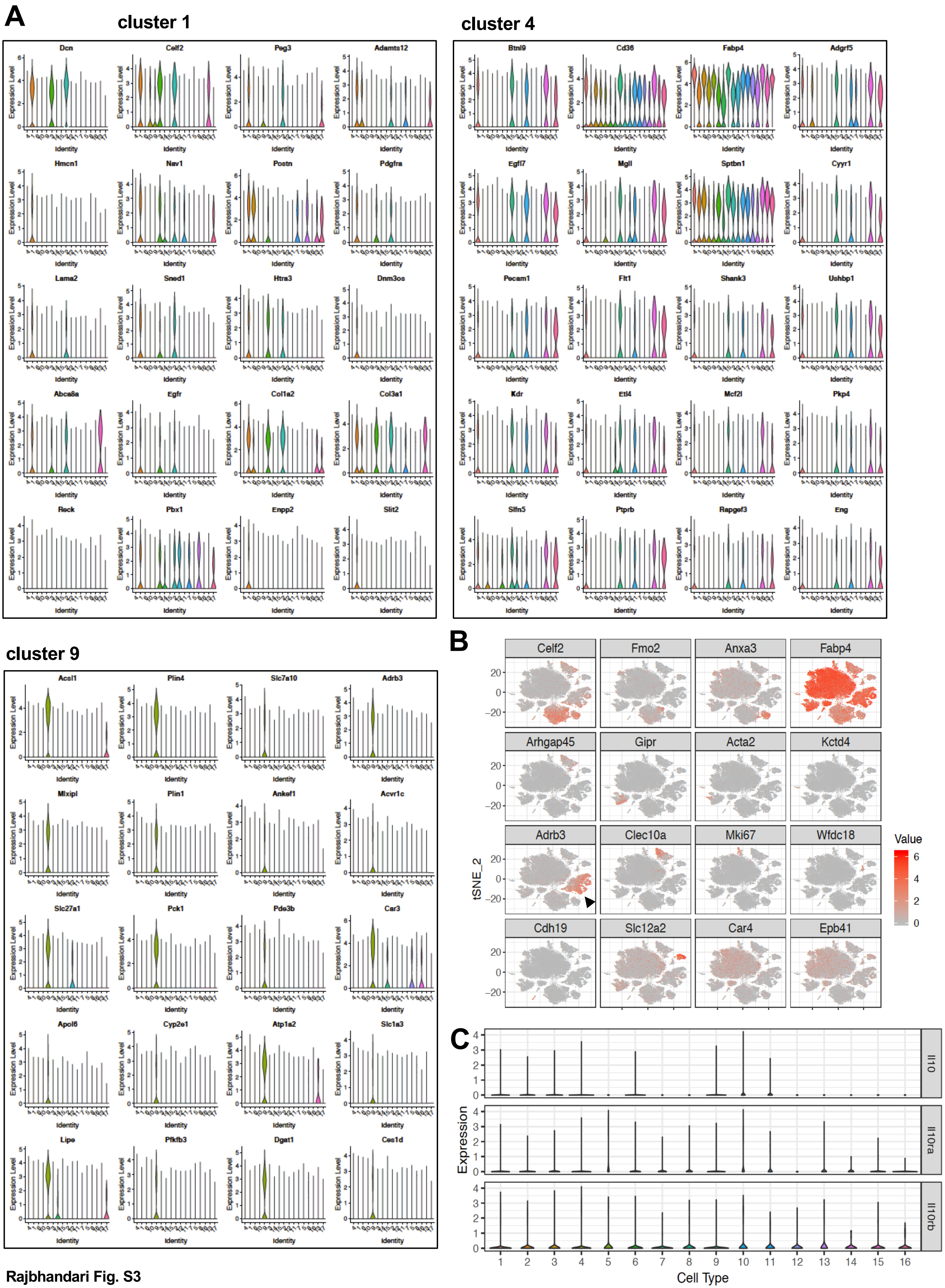


Rajbhandari Fig. S3

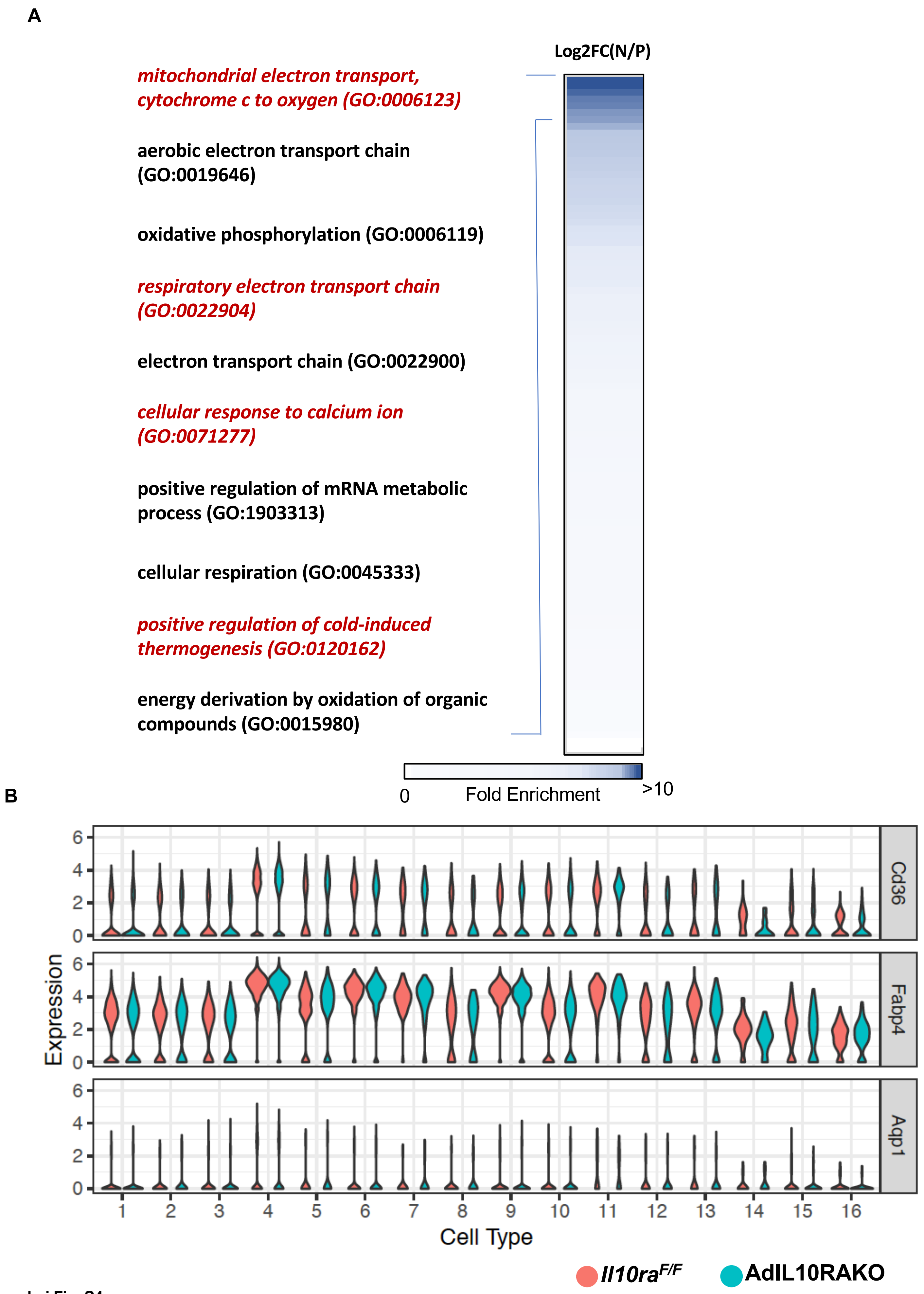


Rajbhandari Fig. S4
